# CV.eDNA: A hybrid approach to invertebrate biomonitoring using computer vision and DNA metabarcoding

**DOI:** 10.1101/2024.09.02.610558

**Authors:** Jarrett D. Blair, Michael D. Weiser, Cameron Siler, Michael Kaspari, Sierra N. Smith, Jessica F. McLaughlin, Katie E. Marshall

## Abstract

1. Automated invertebrate classification using computer vision has shown significant potential to improve specimen processing efficiency. However, challenges such as invertebrate diversity and morphological similarity among taxa can make it difficult to infer fine-scale taxonomic classifications using computer vision. As a result, many invertebrate computer vision models are forced to make classifications at coarser levels, such as at family or order.
2. Here we propose a novel modular method to combine computer vision and bulk DNA metabarcoding specimen processing pipelines to improve the accuracy and taxonomic granularity of individual specimen classifications. To improve specimen classification accuracy, our methods use multimodal fusion models that combine image data with DNA-based assemblage data. To refine the taxonomic granularity of the model’s classifications, our methods cross-references the classifications with DNA metabarcoding detections from bulk samples. We demonstrated these methods using a continental-scale, invertebrate bycatch dataset collected by the National Ecological Observatory Network. We also introduce the CV.eDNA R package, which aims to assist practitioners looking to implement our methods.
3. Using our methods, we reached a classification accuracy of 79.6% across the 17 taxa using real DNA assemblage data, and 83.6% when the assemblage data was “error-free”, resulting in a 2.2% and 6.2% increase in accuracy when compared to a model trained using only images. After cross-referencing with the DNA metabarcoding detections, we improved taxonomic granularity in up to 72.2% of classifications, with up to 5.7% reaching species-level.
4. By providing computer vision models with coincident DNA assemblage data, and refining individual classifications using DNA metabarcoding detections, our methods the potential to greatly expand the capabilities of biological computer vision classifiers. Our methods allow computer vision classifiers to infer taxonomically fine-grained classifications when it would otherwise be difficult or impossible due to challenges of morphologic similarity or data scarcity. These methods are not limited to terrestrial invertebrates and could be applied in any instance where image and DNA metabarcoding data are concurrently collected.

## 1. Introduction

Computer vision has the potential to transform invertebrate ecology by automating estimations of invertebrate abundance, biomass, and diversity (Høye *et al*., 2021; Schneider *et al*., 2022; Blair *et al*., 2024). However, accurately classifying invertebrate species using computer vision is challenging. This is partly due to the sheer diversity of invertebrates, as there are an estimated 7.5 million (∼1.5 million named) terrestrial invertebrates species globally (Stork, 2018). This has led most invertebrate classification models to opt for coarser taxonomic granularity (e.g. order-level instead of species-level classifications) with relatively few unique classification groups (usually <50; (Ärje *et al*., 2020; Blair *et al*., 2022; Schneider *et al*., 2022). However, ecology studies can involve hundreds or thousands of species, which poses a challenge for simpler machine vision techniques.

One way computer vision models have overcome the challenge of handling many thousands or millions of classification labels is by including additional data modalities such as contextual metadata (e.g. collection location) in computer vision models. The mobile app iNaturalist uses this spatiotemporal data in combination with user-submitted photos to classify nearly 80,000 taxa across the tree of life (Leary *et al*., 2023). Other studies have also found substantial improvements to classification accuracy with multimodal models that include both metadata and images (Berg *et al*., 2014; Terry, Roy and August, 2020; Blair *et al*., 2022). However, despite the potential improvements in accuracy, there are several pitfalls to consider when including spatiotemporal metadata in a computer vision model. For one, spatiotemporal metadata is a lagging indicator of species habitat occupancy (i.e. the presence or absence of a species at a given place and time), and as such it is susceptible to data drift over time (Friedland, 2024). That is, spatiotemporal distributions of taxa change over time, but computer vision models can only learn from past data. Unless a computer vision model is updated frequently with more recent data, the species range distributions it has learned may quickly become outdated. Finally, when dealing with many machine learning classes, spatiotemporal metadata does not solve the challenge of gathering enough training data to sufficiently train a computer vision model (Beery *et al*., 2020). In short, studies that incorporate spatiotemporal metadata have shown that supplemental, non-visual data can improve ecological computer vision models, but spatiotemporal metadata itself has several potential drawbacks. In this study, we leverage an alternative data stream that does not pose the same challenges associated with spatiotemporal metadata: DNA metabarcoding.

DNA metabarcoding is an established tool in ecological research that allows for multiple species to be identified from a single sample using high-throughput sequencing (Taberlet *et al*., 2012; Liu *et al*., 2020). Using this method, DNA can be collected from the environment (eDNA) or from preservative media (e.g. ethanol in insect bycatch samples), sequenced, and then used to infer ecological metrics such as species richness and community composition (Marquina *et al*., 2019; Weiser *et al*., 2022). Due to its improved cost-effectiveness, DNA metabarcoding is becoming more frequently used in large-scale studies where traditional morphological identification techniques cannot keep up financially or logistically (Liu et al., 2020). However, despite being an excellent tool for detecting occurrence at fine taxonomic granularity (even below species-level; Stewart and Taylor, 2020), DNA metabarcoding cannot be used to reliably estimate species abundance or biomass (Lamb *et al*., 2019). Instead, eDNA metabarcoding is more suitable for binary presence/absence detections of species. Additionally, while DNA metabarcoding is generally reliable for taxonomic identifications, it is not exempt from false positive and false negative detections (Guillera-Arroita *et al*., 2017). Some examples of how this may occur include DNA contamination and primer mis-priming (false-positives), or DNA degradation and insufficient sampling effort (false-negatives) (Guillera-Arroita *et al*., 2017; Liu *et al*., 2020). Therefore, while DNA metabarcoding offers considerable advantages for biodiversity assessment (e.g., species inventories, species richness) its limitations often necessitate the use of complementary indicators such as visual observations for other metrics (e.g., abundance, biomass) (Schneider et al., 2022).

Given DNA metabarcoding’s ability to produce reliable fine-scale community composition data, and computer vision’s ability to measure abundance and biomass at coarse taxonomic granularity, several studies have called for a synergistic classification pipeline that takes advantage of the strengths of each tool (Schneider *et al*., 2022; Sys *et al*., 2022; Badirli *et al*., 2023). In theory, such a pipeline could leverage DNA metabarcoding’s fine taxonomic granularity against computer vision’s ability to infer specimen-level characteristics (identity, morphology, etc.) to make ecological inferences that would not be possible using either data stream on their own. DNA might also be a favourable alternative to spatiotemporal metadata, as it is a more direct and coincident indicator of species habitat occupancy, likely making it more resistant to data drift over time (Taberlet *et al*., 2018). Despite the potential benefits of multimodal image-DNA classification models for ecological research, few studies have explored this approach. Additionally, proposed hybrid classification pipelines either leave the DNA and image data streams separate (Sys *et al*., 2022), or sequence specimens individually, and thus do not take advantage of metabarcoding’s ability to process bulk samples (Badirli *et al*., 2023; Gong *et al*., 2024).

Here we present a novel modular method for classifying invertebrate taxa that integrates DNA metabarcoding and computer vision. The objective of this hybrid approach is to improve the accuracy and taxonomic granularity of computer vision classifications by adding concurrent community assemblage data derived from DNA metabarcoding into a bulk specimen classification pipeline (Figure 1). The combination of DNA and image data occurs twice throughout the pipeline: first during classification inference in the computer vision model, and then again as a post-processing step for the model’s classifications. While developing this approach, we ask two primary questions: (1) How does error in DNA metabarcoding data affect the accuracy of the computer vision classification model? (2) What are the strengths and limitations of different classification granularity refinement methods? In addition to the case study we present here, we have also developed a GitHub repository to allow our methods to easily be adapted to other study systems (Blair, 2024). The repository introduces the CV.eDNA R package, which contains functions that assist with the implementation of our methods. The repository also includes demonstrative vignettes that walk through the data preparation, model training, and classification granularity refinement steps.

**Figure 1:**
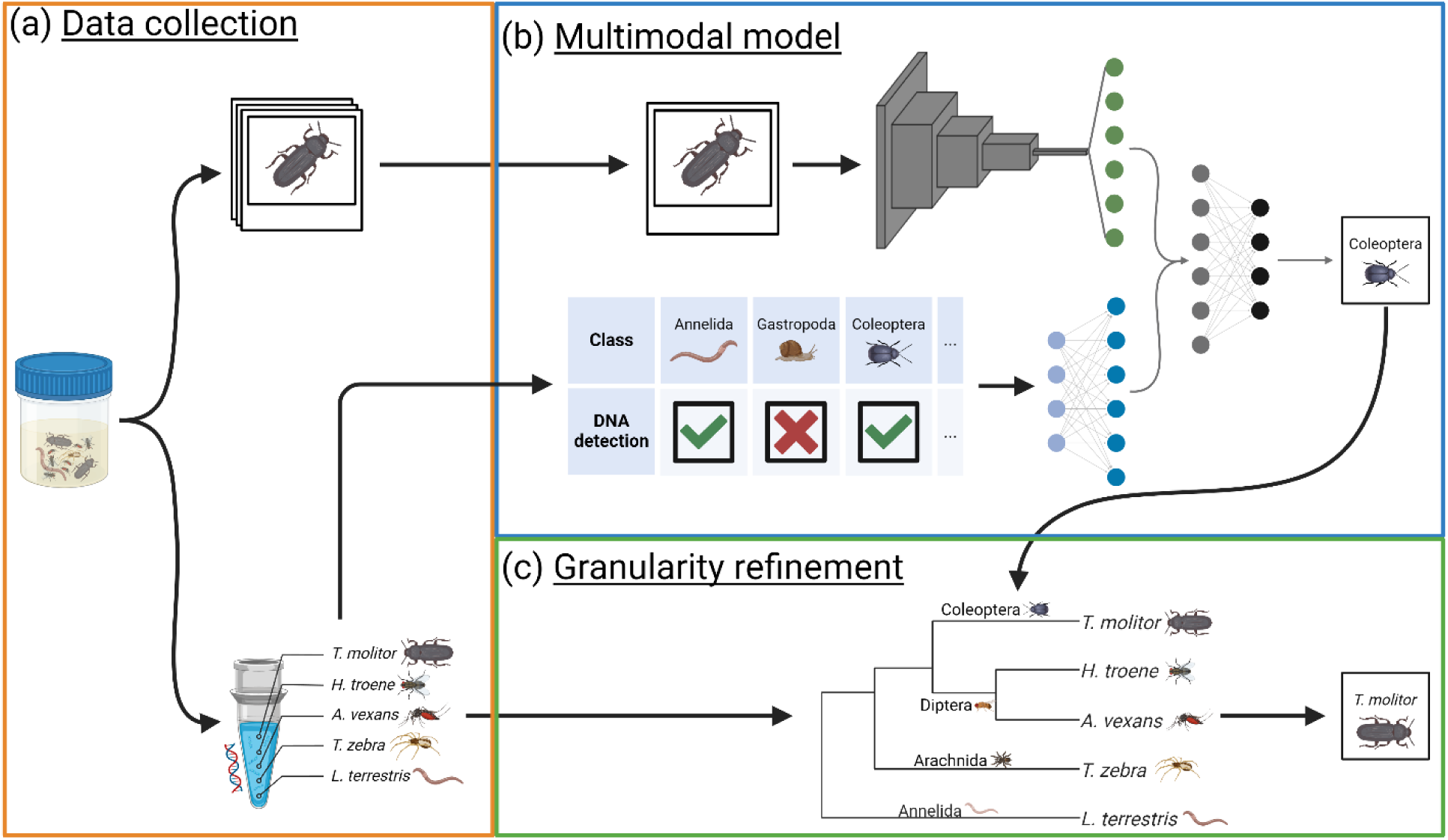
An overview of our methods for combining computer vision and DNA metabarcoding to improve the accuracy and taxonomic granularity of classifications. (a) Images and DNA metabarcoding data are collected concurrently from bulk samples. (b) Images and DNA assemblage data are used as input for a multimodal classification model. The features of the image input are extracted using a convolutional neural network. The DNA assemblage data provides presence/absence information for the model’s known classes and is input as a binary vector into a dense neural network. The image features and the DNA features are concatenated and passed through one more dense layer before final classification in the softmax layer. The visual proportions of each layer have been simplified to ease interpretation and are not meant to be interpreted as 1:1 representations of the exact layer sizes. (c) By interpreting the DNA metabarcoding detections hierarchically and cross-referencing them with the model’s classifications, the taxonomic granularity of the classifications can be refined.

## 2. Methods

### 2.1 Data collection

#### 2.1.1 Specimen collection

Each year, the National Ecological Observatory Network (NEON) performs standardized pitfall trap array sampling across the United States, including Alaska, Hawaii, and Puerto Rico (Hoekman et al. 2017). The focal taxon of the pitfall trap array project are ground beetles (Coleoptera: Carabidae), which are collected, identified, and counted by NEON staff members once every two weeks during the growing season (defined as “the weeks when average minimum temperatures exceed 4 ℃ for 10 days and ending when temperatures remain below 4 ℃ for the same period”, Kaspari et al., 2022). The remaining pitfall trap contents are set aside as ‘Invertebrate Bycatch’ and archived in 95% ethanol-filled 50 mL centrifuge tubes. Hereon, a single collection period from a pitfall trap plot is referred to as a “sampling event”.

The invertebrate bycatch specimens used in this research were taken from 56 NEON trap plots from 27 sites (usually two plots per site; Figure S.1, S.2). Generally, we used three sampling events per plot, selected at the beginning, middle, and end of each site’s growing season. This resulted in a total of 150 sampling events. All sampling events used here were collected in 2016 and processed in 2019. The focus of this project was to classify the invertebrate bycatch, so ground beetles and non-invertebrate specimens were not considered.

#### 2.1.2 Imaging

The contents of each 50mL centrifuge tube were spread out across a 20.32 cm ✖ 30.48 cm (8” ✖ 12”) white ceramic tile and photographed at a resolution of 729 pixels per mm^2^, as described by Weiser et al., 2021 (Figure S.3). Using the FIJI implementation of ImageJ (Schindelin et al., 2012), each specimen was detected and cropped to its bounding box to produce a final image.

#### 2.1.3 DNA extraction and metabarcoding

The DNA metabarcoding data used in this study was collected for Weiser et al., 2022, which used the same sampling events described in Section 2.1.1. In brief, DNA metabarcoding was conducted on a per-tube basis (Figure S.1). Ethanol from each falcon tube was filtered individually (i.e., one filter per tube) and DNA was extracted from the filters using established protocols (Weiser et al., 2022). The cytochrome c oxidase I (COI) barcode region (141-254 base pairs) was then amplified using a two-step polymerase chain reaction (PCR) protocol and sequenced on an Illumina MiSeq. Three COI primers were used: 157, LCO, and Lep (Rennstam Rubbmark *et al*., 2018, 2018; Hajibabaei *et al*., 2019; Weiser *et al*., 2022). Sequences were clustered into Operational Taxonomic Units (OTUs) and each OTU was assigned a taxonomic classification using NCBI BLASTn (Altschul *et al*., 1990) and Integrated Taxonomic Information System (ITIS) (U.S. Geological Survey, 2013). Only sequences with ≥ 97% similarity between the OTU consensus sequence and the BLASTn search were used. See Weiser et al., 2022 for the full DNA extraction and metabarcoding methods.

In total, across all sampling events, there were 10,212 DNA metabarcoding detections. To align the DNA data with the imaging data, we removed any DNA detections from sampling events not included in the image dataset, as well as duplicate detections (i.e. multiple detections of the same taxon in a single sampling event, for example due to amplification using multiple primers). This yielded a final DNA metabarcoding dataset with 3,361 detections and 1,212 unique taxa, primarily consisting of family (369 detections; 85 unique), genus (468 detections; 183 unique), and species-level (2,471 detections; 922 unique) detections.

### 2.2 Data and labelling

#### 2.2.1 Computer vision class labels

The taxonomic scope of the image and DNA metabarcoding data spanned three invertebrate phyla: Annelida, Arthropoda, and Mollusca. The specimen images were labelled by a single technician to the best of their ability (as described in Blair *et al*., 2022). The final labels used for our study ranged from order to phylum-level. Classes with a taxonomic granularity coarser than order-level but with no subtaxa present in the dataset (e.g. Phylum: Annelida) were included.

Specimens labelled as nested classes at a taxonomic granularity coarser than order-level (Phylum: Arthropoda, Class: Insecta, and Class: Arachnida) were excluded, as these classes were primarily composed of “low quality” specimens (highly degraded, low image quality, partial specimens, etc.) that could not confidently be assigned finer level labels. Classes with fewer than 100 specimens in the dataset were also excluded. This resulted in a final image dataset with a total of 36,998 specimens across 17 machine learning classes (13 orders and one subclass, class, subphylum, and phylum; Figure S.4).

#### 2.2.2 Hierarchical labels

Hierarchical labels created by the ‘refhier’ and ‘longhier’ functions in the CV.eDNA package contain taxonomic information at multiple levels (e.g. phylum to species) and can be assigned to images and sampling events (Table S.1, Table S.2). These labels are used as input for the classification granularity refinement methods (see Section 2.5).

In our case study, our image-based hierarchical labels contained taxonomic information at six levels from phylum to order-level for individual specimens (Table S.1). Our DNA-based hierarchical labels contained information at 13 levels from phylum to species for all DNA metabarcoding detections in each sampling event (Table S.2). The levels in the DNA-based hierarchical labels were phylum, subphylum, class, subclass, superorder, order, suborder, infraorder, superfamily, family, subfamily, genus, and species.

#### 2.2.3 Binary assemblage data

The ‘get_assemblagè function in the CV.eDNA package generates binary assemblage data for sampling events. This assemblage data can then be used as class priors or input features for classification models (see Section 2.4). To generate this assemblage data, the function receives DNA metabarcoding data or ground truth image metadata as input and outputs an *n*-element long binary vector for each sampling event, where *n* is the number of classes known by the computer vision model. If a given class is detected in a sampling event, its corresponding element is assigned a score of 1, whereas it is assigned a score of 0 if it was not detected.

For our case study we generated two sets of binary assemblage data containing our 17 known classes: one using detections from the image labels and one using detections from the DNA class labels.

### 2.3 Training and testing data split

Quasi-replication occurs in machine learning datasets when the same or very similar data occur in both the training and testing datasets. This violates the assumption of independence between training and testing data and should be avoided to make valid inferences on the test data. The DNA-based assemblage data presented a quasi-replication risk, as specimens from the same sampling event would have the same assemblage data. To avoid quasi-replication, we split the training and testing such that all specimens from a given sampling event were only included in either the training or testing data. We set our target training:testing ratio to 85:15, and we randomly added sampling events to the test dataset until the test dataset contained >15% of the total number of specimens. The final train:test split was 31,381: 5,617 specimens and 122:28 sampling events.

### 2.4 Classification models

The objective of all the classification models was to accurately classify the class labels of individual specimen photos. Classification masks (Section 2.4.2) and multimodal fusion approaches (Section 2.4.3) were used to assess how classification accuracy changes when DNA-based assemblage data was added to the specimen classification pipeline. Both the classification masks and multimodal fusion models included “oracle” experiments, which used image-based assemblage data to simulate the performance of these methods under optimal conditions. We trained each classification model until the test dataset loss had not improved for 10 epochs (this does not include the classification masks, which did not go through the training process).

Performance of each classification model was assessed using the original image labels as a ground truth. All code required for running these models can be found in the “Model_Scripts” subdirectory of our GitHub repository (Blair, 2024). All models were trained using an AMD Ryzen 7 5800X CPU, an NVIDIA GeForce RTX 4070 Ti GPU and 32 GB of RAM.

#### 2.4.1 Baseline model

To evaluate model performance in the absence of DNA-based assemblage data, we trained a ResNet-50 (He et al., 2016) as a baseline model using only image data. The model was pre-trained using the ImageNet weights from He et al. (2016), and then fine-tuned using the NEON invertebrate bycatch image data. The ImageNet classification layer was removed and replaced with a new classification layer for our 17 classes. A batch normalization and dropout step were also implemented before the final classification layer. The model was trained using the Adam optimizer. The baseline model took 45 seconds per epoch to train, and 7 seconds to run inference on the test dataset.

#### 2.4.2 Classification masks

If *a priori* probabilities for classes in a classification model are known, the outputs of the model can be adjusted according to those probabilities to improve classification accuracy (Saerens, Latinne and Decaestecker, 2002). We implement two variations of this technique which we call “Naïve masks” (Section 2.4.2.1) and “Weighted Masks” (Section 2.4.2.2), whereby the class priors are derived from a sample’s DNA metabarcoding assemblage data. To apply classification masks to our case study, each test dataset specimen’s classification probabilities (i.e. softmax layer values) from the baseline model were multiplied by their sampling event’s classification mask values. The class with the highest classification probability after applying the mask was used as the final classification.

##### 2.4.2.1 Naïve mask

When using a naïve mask, the softmax layer values for a given specimen are multiplied by the specimen’s sampling event’s binary DNA metabarcoding assemblage data (generated using the ‘get_assemblagè function). Thus, any classes not detected by the DNA metabarcoding in a given sampling event have their respective softmax values set to 0, whereas the remaining classes are unaffected. Naïve masks essentially operate as a “hard filter”, where only classes detected by the DNA metabarcoding can be classified by the model.

##### 2.4.2.2 Weighted mask

“Hard” masks like the naïve mask, which set the softmax values of undetected classes to 0, assume the DNA metabarcoding data is error-free. However, in reality, DNA metabarcoding can have false positive and/or false negative detections (Taberlet *et al*., 2018). A weighted mask is a “softer” version of a naïve mask that allows classes not detected by the DNA metabarcoding to still be classified. The weights for a weighted mask can be generated using the ‘get_weights’ function of the CV.eDNA package, and are calculated using the metabarcoding’s true positive rate (precision) and false negative rate (1 - recall) for each class. The DNA metabarcoding precision and recall are calculated by comparing the DNA-based assemblage data to ground-truth assemblage data (e.g. manually assigned image-based assemblage data). The weight that is assigned to any given class in a sample is determined by whether or not the class was detected by the DNA metabarcoding. Classes which have their DNA detected are assigned their metabarcoding precision value, while classes which do not have their DNA detected are assigned their 1 – recall value.

##### 2.4.2.3 Oracle mask

As an oracle experiment to simulate a scenario where the DNA detections were in perfect alignment with the image-based detections, we created a naive classification mask using the image-based assemblage data. This mask was then applied to the classification output of the baseline model. We did this to provide an upper-bound for the classification mask accuracy, and to understand how DNA detection accuracy impacts specimen classification accuracy when using classification masks.

#### 2.4.3 Multimodal fusion

In deep learning, multimodal models can receive data from multiple modalities as input to inform their classifications (Ramachandram and Taylor, 2017). For example, previous studies have described multimodal classification models that receive images and raw DNA barcode sequence data for individual specimens (Badirli *et al*., 2023; Gong *et al*., 2024). However, no previous studies have described a multimodal model that combines specimen images with DNA metabarcoding data from bulk samples. We achieve this by inputting DNA metabarcoding data as tabular binary assemblage data, which is then combined with specimen image features using intermediate fusion (Figure 1b). Intermediate fusion is an approach to building multimodal models where data from each modality is input separately. The features from each input are then extracted and concatenated prior to classification (Boulahia *et al*., 2021). This allows the model to contextualize a specimen’s image features with class presence-absence data from the specimen’s sample. However, similar to the weighted mask approach (Section 2.4.2.2), the DNA metabarcoding data does not act as a “hard filter”, and the model can still make classifications not detected by DNA metabarcoding. Additionally, because the assemblage data contains all class detections for a given sample, the model can learn patterns of class co-occurrence to inform its classifications.

In our case study, we paired our DNA metabarcoding assemblage data with individual specimens based on their sampling event (the same approach as the classification masks described in Section 2.4.2). After being input, features of the assemblage data were extracted using a single fully-connected layer with batch normalization and dropout. Specimen images were fed through the ResNet-50 architecture, which ultimately producing a flat feature layer. The image and assemblage feature layers were then concatenated and passed through another fully-connected layer with batch normalization and dropout before reaching the final classification layer. During training, both sides of the multimodal model were trained simultaneously using the Adam optimizer.

To understand the effect that DNA detection accuracy has on classification performance, we ran three versions of the multimodal model using different types of assemblage data as input: (1) using the DNA-based assemblage data, (2) using the image-based assemblage data, and (3) using ‘zero-filled’ assemblage data (all values in the assemblage data are set to zero). In all three experiments the training and testing datasets used the same assemblage data type (i.e. DNA-based, image-based, or zero-filled). All three experiments used the same overall model architecture as described in Figure 1b. The purpose of the image-based assemblage experiment was to simulate the results of a model where the DNA detections perfectly aligned with the ground truth labels (i.e. an oracle experiment). The purpose of the zero-filled assemblage experiment was to control for differences in model architecture when comparing the multimodal models to models trained without DNA-based assemblage data, as the zero-filled data would provide no informative value to the model. The zero-filled assemblage data had the same dimensions as the other assemblage data (17 values for each sampling event). The multimodal models took 105 seconds per epoch to train, and 16 seconds to run inference on the test dataset.

### 2.5 Refining taxonomic granularity using DNA-based assemblage data

A strength of DNA metabarcoding is its ability to produce species-level detections. The ‘modelbias’ and ‘dnabias’ functions in our CV.eDNA package take advantage of this strength to refine the taxonomic granularity computer vision model classifications by cross-referencing them with the DNA metabarcoding detections (Figure 2). When using either function in cases where the model classifications and DNA metabarcoding detections agree on the presence of a class, the granularity of the classification improves until the number of subtaxa detected by the DNA metabarcoding was greater than 1 or the granularity reached species-level (Figure 2a,b). The functions differ in cases where the model classifications and DNA detections disagree on the presence of a class classified by the model. The model-biased method of the ‘modelbias’ function is simple. In cases where the model and DNA metabarcoding disagree, the model classification remains unchanged (Figure 2c). When using the DNA-biased method of the ‘dnabias’ function, the hierarchical label of the classified specimen is cross referenced with the DNA metabarcoding hierarchical labels of the sampling event (Figure 2d, Table S.1, Table S.2). Starting from the original classification level, the granularity of the label is coarsened until an agreement between the model and DNA metabarcoding is reached. The taxonomic name at this level becomes an intermediate label, which is refined until the number of subtaxa detected by the DNA metabarcoding is greater than 1 or the granularity reached species-level.

**Figure 2:**
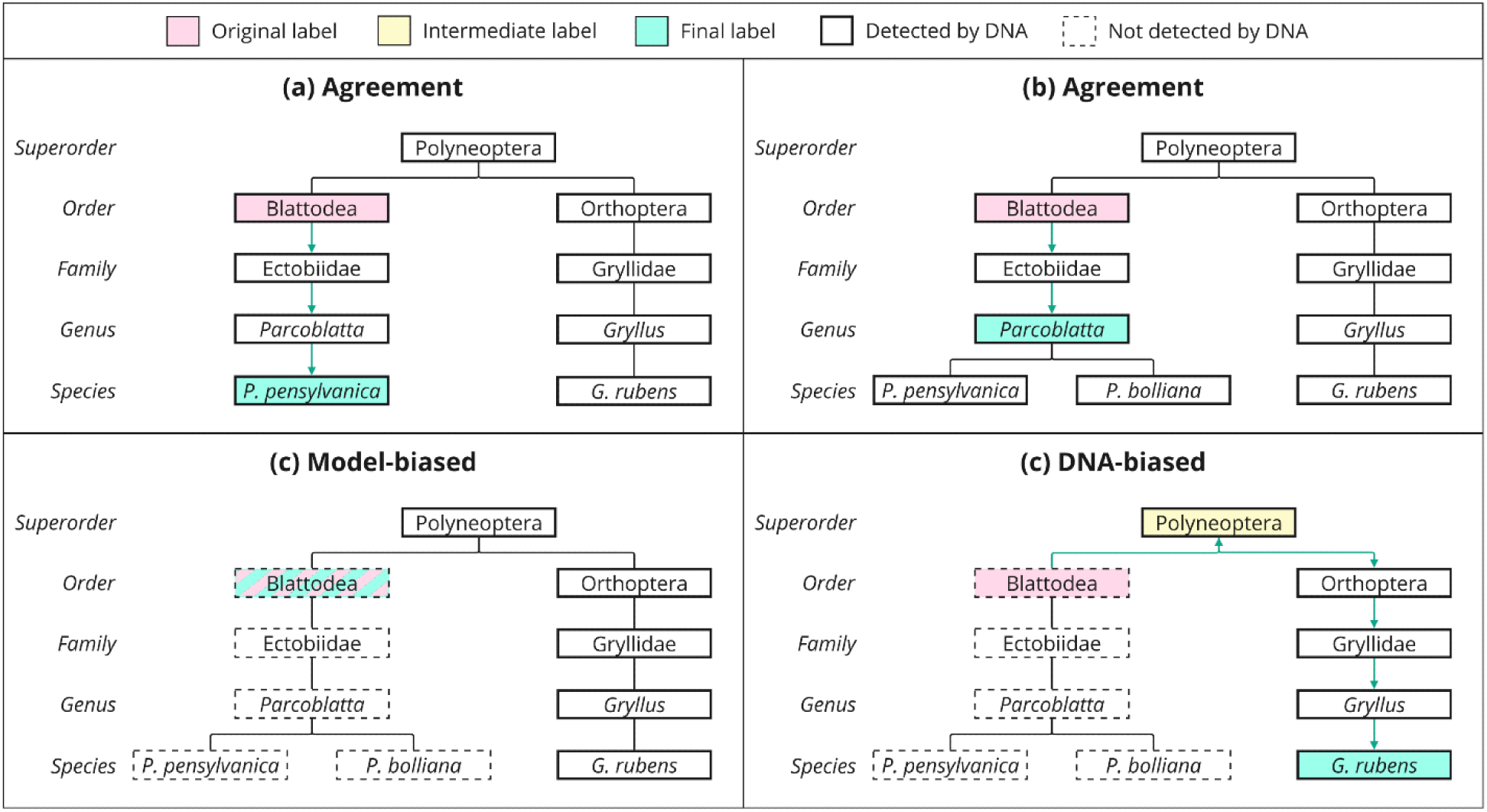
Four methods of changing a label’s granularity using DNA detections. (a,b) When the classification label and DNA detections are in agreement about the presence of a class, granularity will be refined until the number of subtaxa detected by the DNA metabarcoding is > 1 or the classification reaches species level. (c) Under the model-biased approach, when the classification label and DNA metabarcoding do not agree on the presence of a class, the classification label remains unchanged. (d) Under the DNA-biased approach, when the classification label and DNA metabarcoding do not agree on the presence of a class, the granularity of the classification label is coarsened until a DNA detection for the class is found (“intermediate label”). The granularity of the intermediate label is then refined using the same rules as (a,b).

To test the effectiveness of each method, we applied the model-biased and DNA-biased methods to the classifications of our DNA multimodal fusion model. A vignette for these methods can be found on our GitHub repository (Blair, 2024).

## 3. Results

### 3.1 DNA metabarcoding precision and recall

To generate the weights for the weighted mask, we calculated the DNA-based assemblage data’s precision and recall for each class using the image-based assemblage data as the ground truth. This calculation only included sampling events from the training dataset. Across the 17 predicted classes, recall ranged from 0.905 to 0.000, with an average recall of 0.570. The weight for a negative detection was 1 - recall, so negative detection weights ranged from 1.000 to 0.095, with an average of 0.430. Two classes (Opilioacarida and Zygentoma) were never detected by the DNA. Across the 15 classes detected by the DNA, the average precision was 0.761. The precision and recall values per class are reported in Table S.3.

### 3.2 Classification accuracy

Compared to the baseline model, the DNA multimodal fusion model improved accuracy by 2.2% (79.6% vs 77.4%; Table 1). However, the DNA multimodal fusion model’s accuracy was 1.0% lower than the zero-filled model’s accuracy (80.6%). This suggests that some of the performance improvements seen in the multimodal fusion models could come from changes to the model architecture. The naïve classification mask recorded the lowest classification metrics, with an accuracy 22.6% below the baseline model (54.8%), and a top-3 accuracy of only 64.0%.

**Table 1:** Performance metrics for experiments trained using DNA-based assemblage data, other than the baseline model, which was trained only using images, and the ‘zero-filled’ experiment, which replaced all assemblage data values with zero to control for the impact of model architecture. Underlined scores indicate they are the highest for a given metric.

| Experiment | Accuracy | Balanced Accuracy | Top-3 Accuracy |
| --- | --- | --- | --- |
| Baseline | 0.774 | 0.674 | 0.952 |
| Naive mask | 0.548 | 0.509 | 0.640 |
| Weighted mask | 0.764 | 0.666 | 0.950 |
| Multimodal fusion | 0.796 | <u>0.680</u> | <u>0.957</u> |
| Multimodal fusion (zero-filled) | <u>0.806</u> | 0.670 | 0.952 |

Our oracle experiments with image-based assemblage data performed better, with the multimodal fusion model reaching an average accuracy of 83.6% and a balanced accuracy (i.e. macro-averaged recall) of 0.713 (Table 2). The image-based naive mask accuracy was 25.8% better than the DNA-based naive mask, and 3.2% better than the baseline model. It also had the highest top-3 accuracy across all experiments at 93.2%. Training dataset accuracy results for all non-mask experiments are reported in Table S.4.

**Table 2:** Performance metrics for the oracle experiments using image-based assemblage data. These experiments used binary assemblage data taken from the ground truth specimen labels. Underlined scores indicate they are the highest for a given metric.

| Experiment | Accuracy | Balanced Accuracy | Top-3 Accuracy |
| --- | --- | --- | --- |
| Mask | 0.806 | 0.690 | 0.959 |
| Multimodal fusion | <u>0.836</u> | <u>0.713</u> | <u>0.963</u> |

### 3.3 Taxonomic granularity

Of the DNA multimodal fusion model classifications, 68.2% (3833/5617) were present in their corresponding sampling event’s DNA assemblage data. Nonetheless, both approaches for handling disagreements between the DNA and classification model (model-biased or DNA-biased) improved the average taxonomic granularity of the classifications (Figure 3). Despite starting with no classifications finer than order-level, the DNA-biased approach resulted in 5.7% of classifications improving to species-level and the model-biased approach resulted in 5.1% of the classifications improving to species-level. The DNA-biased approach was more effective overall at refining the granularity of classifications, with 72.2% of classifications becoming finer than their original classification, and 43.1% reaching at least family-level. In the model-biased approach, 51.7% of classifications improved their granularity, and 31.0% reached family-level or lower. When looking exclusively at classifications where the DNA assemblage data and multimodal fusion model classifications agreed on the presence of a class, 7.5% reached species level, 45.4% reached family-level or lower, and 75.7% became finer than their original classifications. Both the DNA-biased and model-biased approaches produced 89 unique final labels (Table S.5).

**Figure 3:**
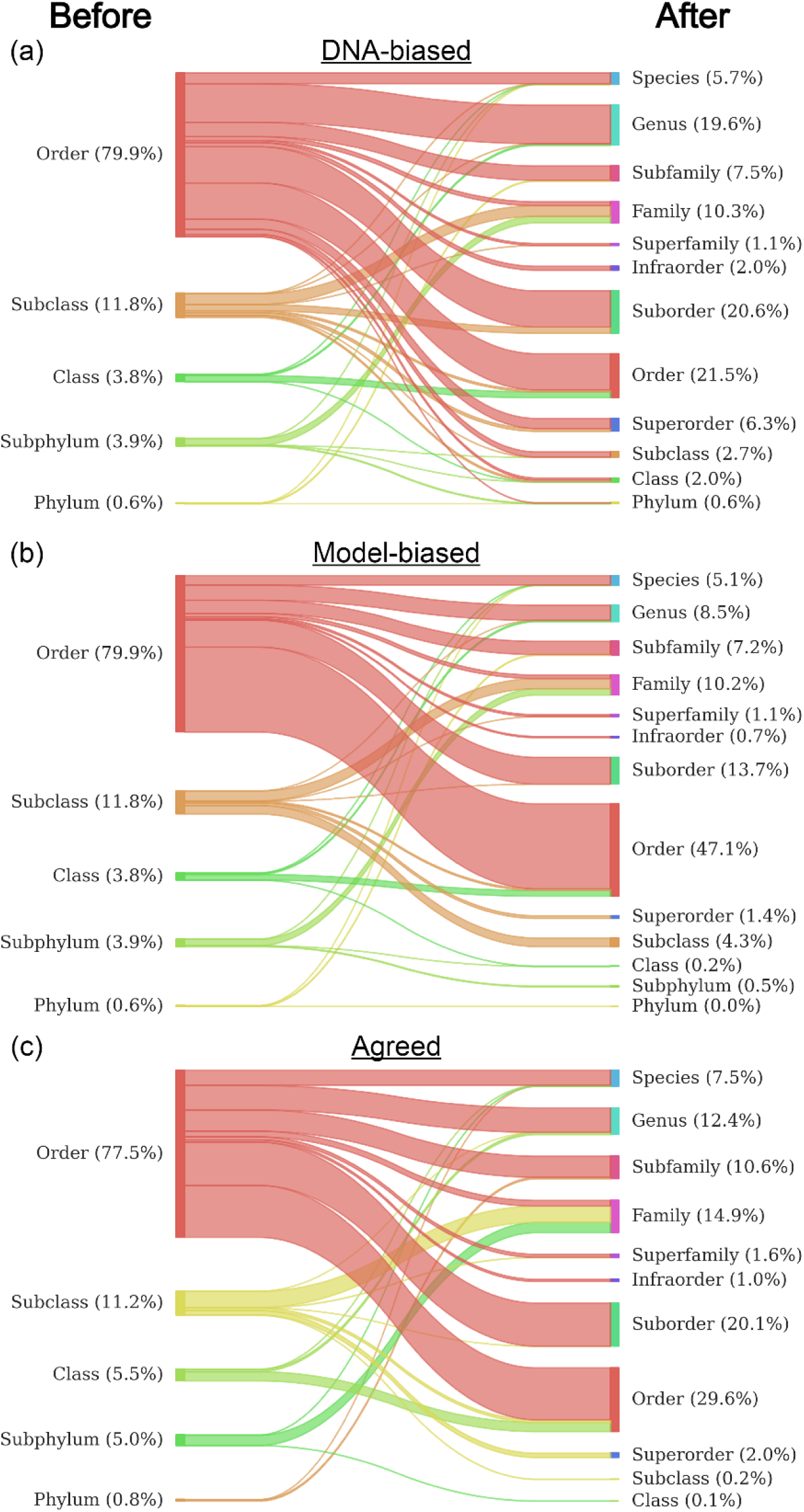
Sankey diagrams showing the change in taxonomic granularity before (left) and after (right) cross-referencing labels with the DNA detections. (a) DNA-biased approach. (b) Model-biased approach. (c) Results when the model classification and DNA detections agree on the presence of the labelled class.

## 4 Discussion

Here we show that combining concurrent DNA metabarcoding assemblage data with computer vision can improve the accuracy and taxonomic granularity of computer vision classifications. Unlike most classification pipeline enhancements which only focus on improving the ability to classify classes on which it was trained (“known classes”), this approach adds the ability for the pipeline to infer classifications beyond the model’s usual taxonomic scope (“unknown classes”). Our methods could be applied to any study system where images and DNA metabarcoding data are collected concurrently, provided that the practitioners have labelled data to train their own classification model. To thoroughly explore the benefits and implications of our hybrid approach, we focused on the following two research questions.

### 4.1 How does DNA metabarcoding accuracy affect specimen classification accuracy?

The effect of metabarcoding accuracy on classification accuracy differed between the classification masks and the multimodal fusion models, illuminating a key difference between them: classification masks (as used in this study) do not take class co-occurrence into consideration, whereas the multimodal models do. Put another way, in a classification mask the only factor that directly influences the weight given to a class is the presence or absence of the class itself. Conversely, the neural networks of multimodal fusion models—with their fully-connected structure—allow the presence or absence of all classes to holistically influence each class’s classification probability. This allows the model to use patterns of class co-occurrence to inform its classification decisions.

The different mechanisms used by classification masks and multimodal models are best demonstrated in Table 2, where the assemblage data was derived from the ground-truth image labels. In the classification mask, the model’s original classifications were exclusively based on image data, and any classifications that did not match their respective assemblage data were reclassified as the class with the highest softmax score that was present in the specimen’s assemblage. Given that the assemblage data was derived from the ground truth labels, this mask acted as a sieve that filtered out any classes that could not possibly be the correct classification based on the assemblage data. As a result, it could only have a positive impact on accuracy.

However, despite this, the multimodal fusion model still scored higher on all three metrics measured (top-1 accuracy, balanced accuracy, and top-3 accuracy). This implies that the multimodal fusion model was not just using the assemblage data as a filter, but that it provided additional contextual information (such as class co-occurrence or class exclusion) that further improved accuracy. Thus, due to the multimodal fusion model’s ability to holistically evaluate occurrence data, and as illustrated through its superior performance compared to classification masks even under ideal conditions, multimodal fusion models are likely to be preferable in most use cases. This conclusion is reinforced by the results of Table 1, where both naïve and weighted masks showed negative effects on all classification performance metrics when the DNA-based assemblage data contained substantial amounts of error.

### 4.2 What are the strengths and limitations of each granularity refinement method?

Here we proposed two approaches for cross-referencing DNA metabarcoding data to achieve the novel ability of refining the taxonomic granularity of computer vision classifications. The two approaches differ in how they resolve disagreements between the detections of the DNA metabarcoding and computer vision classifications, with the model-biased approach favouring the computer vision classifications, and the DNA-biased approach favouring the DNA detections. As such, each approach has its own set of advantages and limitations.

Through the ability to coarsen granularity before refining it, the DNA-biased method can make classifications outside of the taxonomy of the original classification model (Figure S.5).

Explained another way, the model-biased approach and traditional hierarchical classifiers (e.g. Badirli *et al*., 2023) can only adjust classifications “vertically” (i.e. to supertaxa or subtaxa of the original classifications), but the DNA-biased approach can also adjust classifications “laterally” to out-of-distribution taxa through a combination of classification coarsening and refining.

Classification of out-of-distribution taxa is usually only possible using feature embedding learning methods such as zero-shot learning (Badirli *et al*., 2021). An illustrative example of this comes from the DNA-biased approach’s detection of taxa within the insect order Psocodea (e.g. *Valenzuela flavidus*; Table S.5). As Psocodea was not included as a class in our model, and our model’s finest taxonomic granularity was order-level, Psocodea’s branch of the taxonomy was only accessible through a “lateral” taxonomic adjustment (Figure S.5). As such, it was only detected by the DNA-biased approach, and not the model-biased approach (Table S.5). In theory, this extends the range of possible classifications to the full taxonomic scope of the genetic reference database being used (e.g. GenBank, Barcode of Life, etc.) (Ratnasingham and Hebert, 2007; Sayers *et al*., 2024). In practice, it is likely best to self-impose limits on how much the DNA-biased method can coarsen granularity. In our case we limited ourselves to phylum, as we were only interested in classifications within our three focal phyla.

While it does not have the same potential taxonomic scope of the DNA-biased approach, an advantage of the model-biased approach is that the taxonomic granularity of the final classification cannot be coarser than the original classification. Applied to the DNA multimodal fusion model classifications, 9.9% of all classifications became coarser when the DNA-biased method was used (Figure 3). While the DNA-biased method classified more specimens at family-level or finer (43.1% vs 31.0%), the ability to coarsen granularity resulted in more classifications above order-level (11.7% vs 6.4%).

Even when granularity does not reach species, classifications that match with the DNA-based assemblage data still provide more information than what is typically output from a classification model. This is because we can also see the number and identity of subtaxa that the specimen could be according to the DNA metabarcoding detections. For example, if the DNA metabarcoding detected three species of the cricket genus *Gryllus* in a sample, we could say the label of a specimen that would otherwise be classified simply as “*Gryllus* indet.” is actually one of three possible species of *Gryllus*, as detected by the DNA metabarcoding (e.g. *G. pennsylvanicus*, *G. rubens*, or *G. veletis*). This might also be useful for future developments to these methods, as the number of subtaxa detected by the DNA metabarcoding could be used to inform clustering algorithms that separate the specimens into morphotaxa.

Of course, the granularity of classifications matters little if they are not accurate. A caveat of our study is that we cannot verify the accuracy of granularity-refined classifications, as they are at lower taxonomic levels than our ground truth (human-classified) labels. However, we know that our classification models were more accurate than the DNA assemblage data when compared to our ground-truth labels. Thus, the DNA-biased method of refining granularity will likely add more error to the classifications than the model-biased approach. When deciding between the two methods, this is likely to be a determining factor: does the computer vision model or DNA metabarcoding contain more error?

### 4.3 Caveats and areas for future exploration

In an applied context, we cannot definitively conclude that image-DNA multimodal fusion models as we present here improve specimen classification accuracy. This is primarily due to the high rates of disagreement (or “error”) between our DNA metabarcoding detections and image-based detections. When comparing our three multimodal fusion experiments, the zero-filled experiment had a top-1 accuracy 1.0% higher than the DNA-based assemblage data experiment, but 3.0% less than the oracle experiment. This suggests that in an ideal situation where the DNA-based assemblage data has low amounts of error (i.e. it is more similar to the image-based assemblage data), image-DNA multimodal fusion models will positively impact classification accuracy. However, when the DNA-based assemblage data contains substantial error, differences in performance between the baseline and multimodal fusion models likely arise from changes in the model’s architecture.

Reconciling genetic-based and morphology-based data—the two chief methods for invertebrate biodiversity monitoring—is a pressing need as previous studies have shown that assemblages determined by visual classification usually differ from assemblages determined using DNA metabarcoding. For example, Emmons *et al*. (2023) found that NEON benthic macroinvertebrate samples classified by taxonomists only shared 59% of order-level detections with DNA metabarcoding data derived from homogenized blends of the same samples. Similar results have also been produced in other studies comparing metabarcoding methods to morphologic identification (Remmel *et al*., 2024; Salis *et al*., 2024). Marquina *et al*., (2019) also found that different DNA sampling protocols can produce inconsistent assemblage data, as DNA metabarcoded from ethanol vs homogenized blends of the same samples yielded significantly different assemblage data, with both methods detecting taxa not detected by the other. Despite these challenges, solutions to improve invertebrate detections using DNA metabarcoding are being investigated. Proposed solutions range from changes in sampling methodology (e.g. strategically subsetting bulk samples; (Remmel *et al*., 2024) to improvements in the completeness of publicly available DNA reference databases (Salis *et al*., 2024). For the image- DNA multimodal fusion methods we propose here to be maximally effective, advances will need to be made in DNA metabarcoding methodology to limit false positive and false negative detections.

Beyond DNA metabarcoding accuracy, there are likely other factors that can impact the efficacy of our methods, such as the alpha and beta diversity of the sample assemblages. Lower alpha diversity and higher beta diversity should yield models with greater classification performance. Higher beta diversity would improve classification performance because the composition of the assemblages would be more heterogeneous, which is required for learnable patterns to emerge in the data. For example, our zero-filled assemblage experiment used data that was completely homogenous, as every sampling event had the same assemblage data, thus allowing no learnable patterns to emerge from the assemblage data. Conversely, lower alpha diversity would improve classification performance because more classes to be filtered out by the model. This is partially demonstrated by comparing the results of Blair *et al*. (2020) to the results we present here. In their study, which built classification models for NEON’s carabid beetles, the authors applied classification masks to their models based on the detected ground beetle assemblages at each sampling site. On average, 2.93 out of 25 potential species (11.7%) were detected per site, resulting in an accuracy improvement of 10.9% (84.7% → 95.6%) after applying the classification masks. Comparatively, our image-based assemblages detected an average of 9.09 out of 17 potential classes (53.5%) per sampling event. When compared to the baseline model, this resulted in an accuracy improvement of 3.2% when using the classification mask and 6.2% in the multimodal fusion model (Table 1, Table 2). Sample alpha diversity also has an impact on the efficacy of the classification granularity refinement step. Higher alpha diversity increases the odds that related taxa could present in the same sample, which can force our granularity refinement methods to stop at a coarser taxonomic rank (Figure 2). Thus, sampling methods that increase sample beta diversity (e.g. finer-grain class labels; Terlizzi *et al*., 2009) and reduce sample alpha diversity (e.g. smaller sample size; Chiu, 2023) will likely increase the efficacy of image-DNA metabarcoding multimodal fusion classification pipelines.

### 4.4 Broader applications and implications

In this study, we used assemblage data derived from DNA metabarcoding to improve computer vision classifications of terrestrial invertebrates. Previous studies (Badirli *et al*., 2023; Gong *et al*., 2024) have paired images of specimens with their DNA barcode sequence as input for multimodal classification models. However, this technique cannot be applied to DNA metabarcoding data because samples are metabarcoded in bulk, so the metabarcoded sequences cannot be paired with individual specimens. Our methods overcome this challenge by converting DNA metabarcoding data to binary assemblage data, which can then be input to a multimodal classification model. This advancement has several practical implications, as DNA metabarcoding’s ability to process bulk samples means that it is more time efficient, less expensive, and produces less waste than individual specimen barcoding (Gueuning *et al*., 2019).

Our method’s ability to refine classification granularity, which is typically not possible in computer vision, could improve the feasibility of building broad-scope, fine-grain classification models (e.g. models spanning entire classes or phyla and capable of producing species-level classifications). This typically requires vast amounts of training data, as training examples need to be provided for every species. Using the approach that we present here, classifiers could be trained at coarser taxonomic levels such as order or family and still have the potential to produce species-level classifications. This would decrease the number of classes in the model, and thus data needed to train it, by orders of magnitude. Hence, the synergy between DNA metabarcoding and computer vision outlined in this study paves the way for new possibilities in computer vision classification of taxa, with the potential for improved accuracy and granularity with far less data dependency.

## Author contributions

JDB and KEM conceived the ideas and designed methodology; JDB, MDW, CS, SNS, and JFM collected data and helped refine ideas; JDB analysed the data and led the writing of the manuscript. KEM and MK were principal investigators. All authors contributed critically to the drafts and gave final approval for publication.

## Notes

https://github.com/Jarrett-Blair/CV-DNA-Hybrid

### Competing Interest Statement

The authors have declared no competing interest.

### Summary of Updates

An author contributions section was added.

