## Supplemental materials for "CV.eDNA: A hybrid approach to invertebrate biomonitoring using computer vision and DNA metabarcoding"

Figures

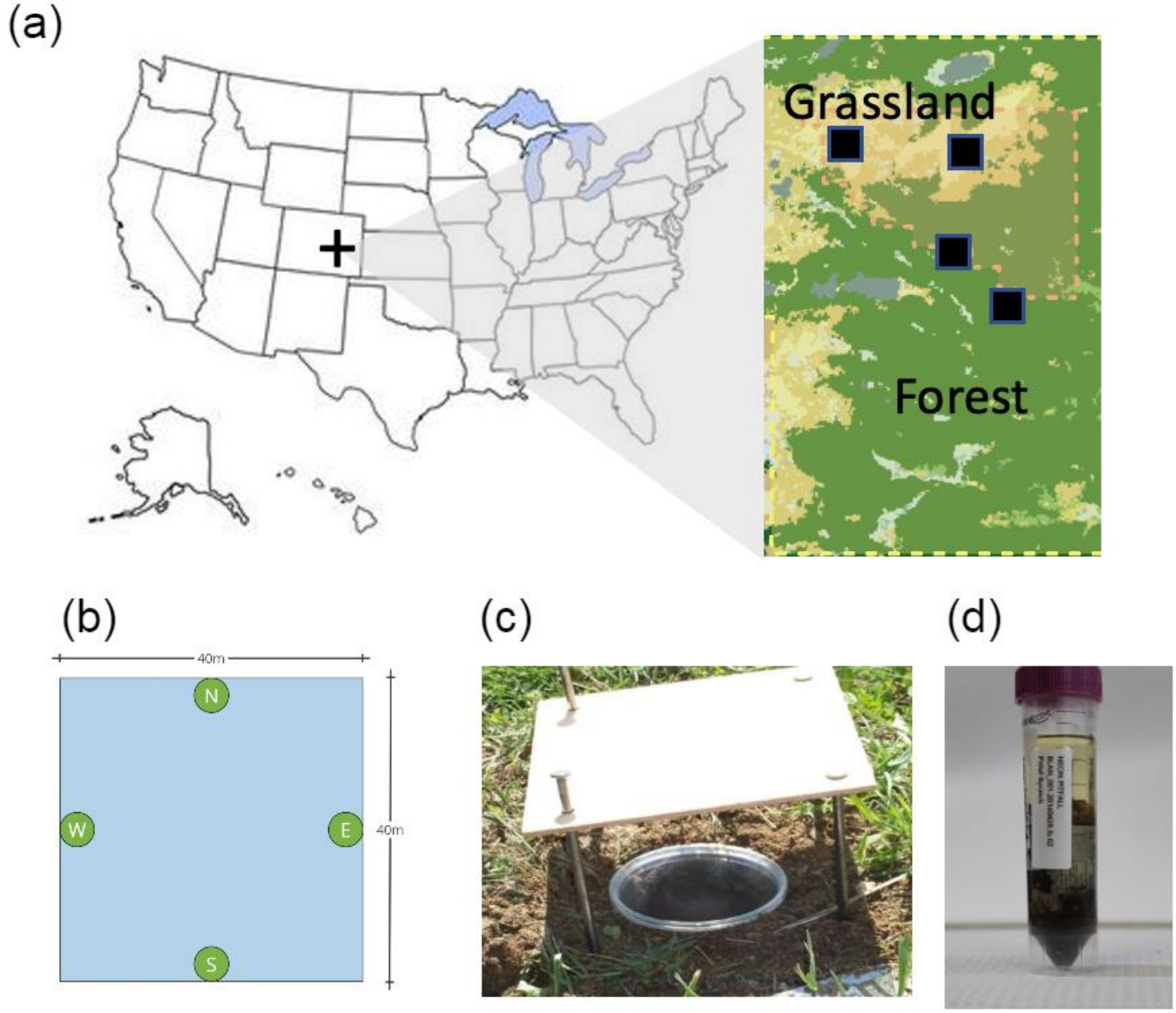


Figure S.1: The National Ecological Observatory Network (NEON) pitfall trap sampling protocol. (a) Sites are distributed across the United States and Puerto Rico. Each site contains an array of sampling plots. This map was originally published in Kaspari et al., 2023 and has been modified for use here. (b) Each sampling plot is 40m × 40m and has four pitfall traps evenly spaced around the perimeter of the plot. This diagram is a simplified recreation of a figure from Hoekman et al., 2017. (c) Pitfall trap (image taken from neonscience.org). (d) Contents of a pitfall trap in a 95% ethanol-filled storage tube.

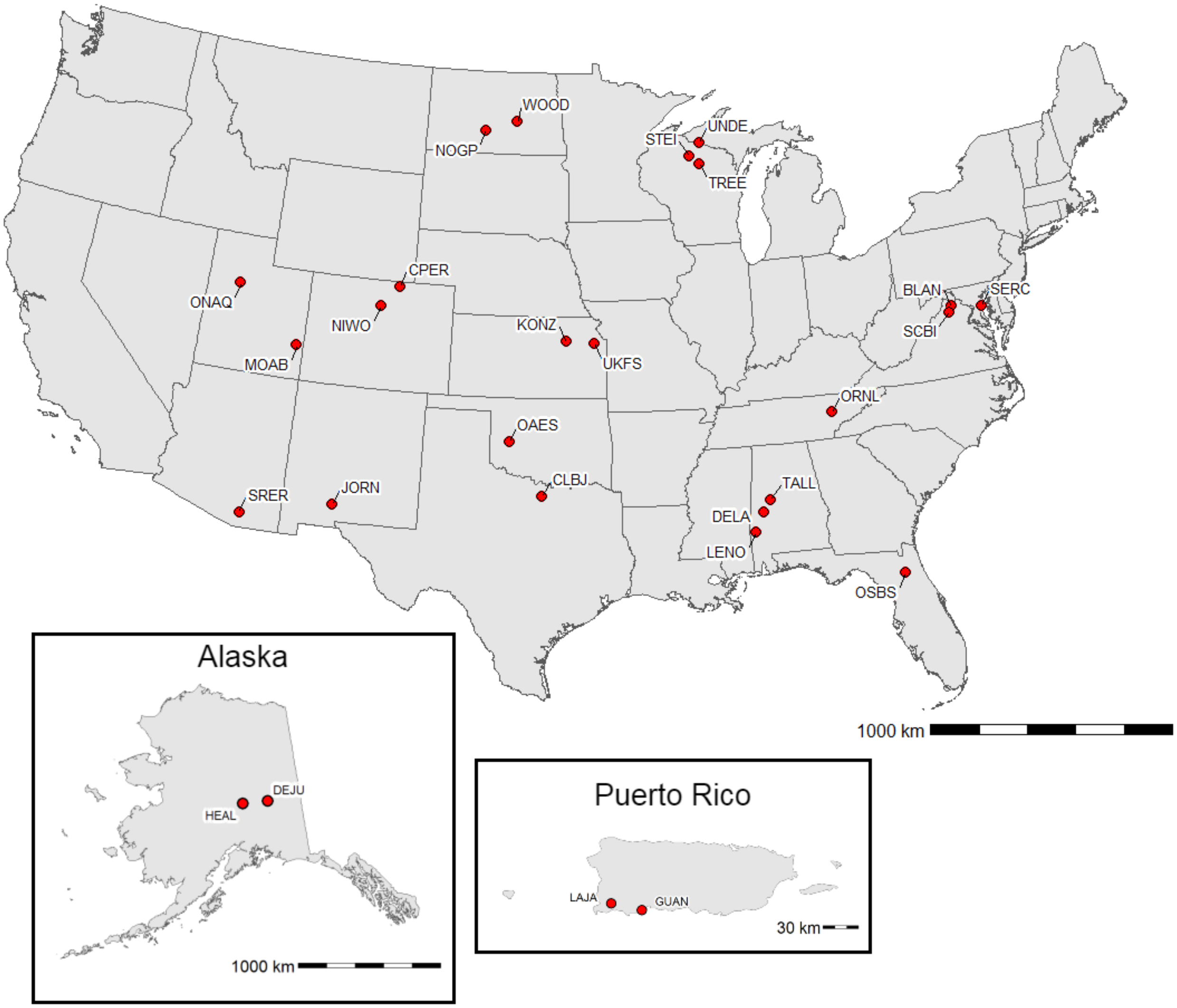


Figure S.2: Map of the 27 NEON sampling sites used in this study. The sites are labelled with their abbreviated names.

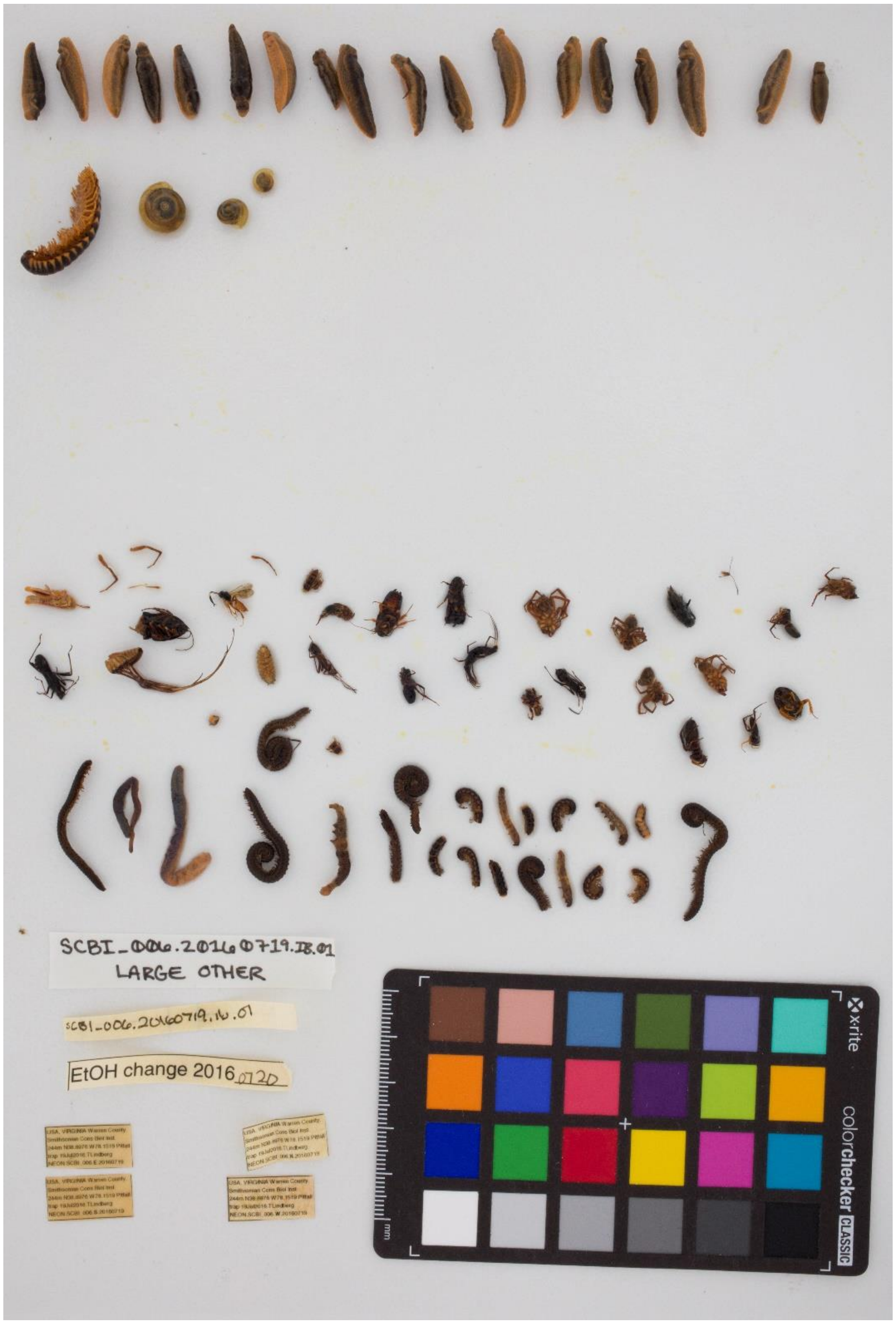


Figure S.3: An example raw, bulk image of National Ecological Observatory Network (NEON) invertebrate bycatch specimens. Specimens were detected and cropped from these images to produce the final images that were used for model training and testing.

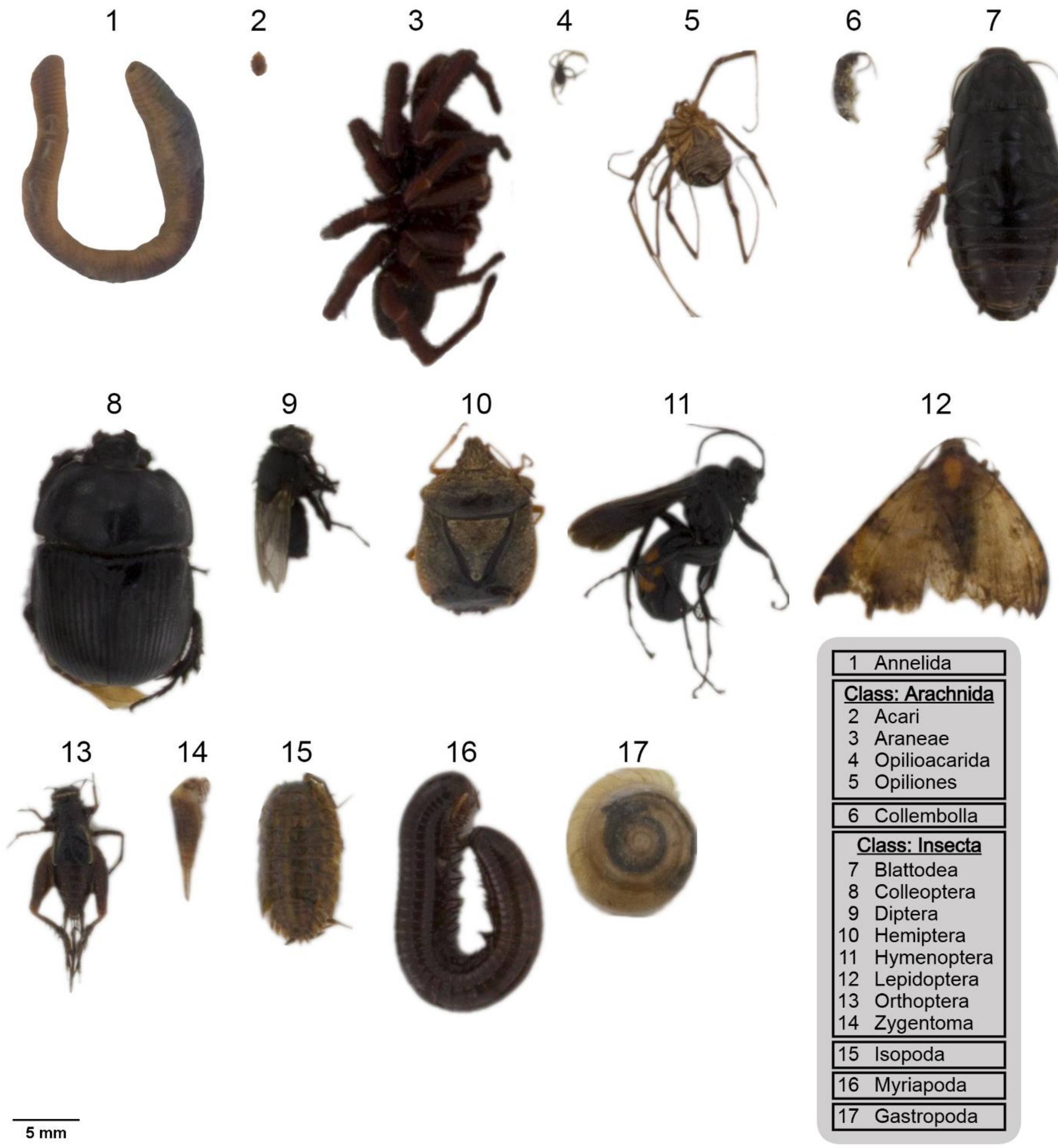


Figure S.4: Collage of photographs of all invertebrate classes used in the training dataset (n = 17). The taxonomic granularity of the classes ranges from order to phylum. Specimens were cropped from their original photographs, and the background was removed. Relative scale of each specimen is conserved.

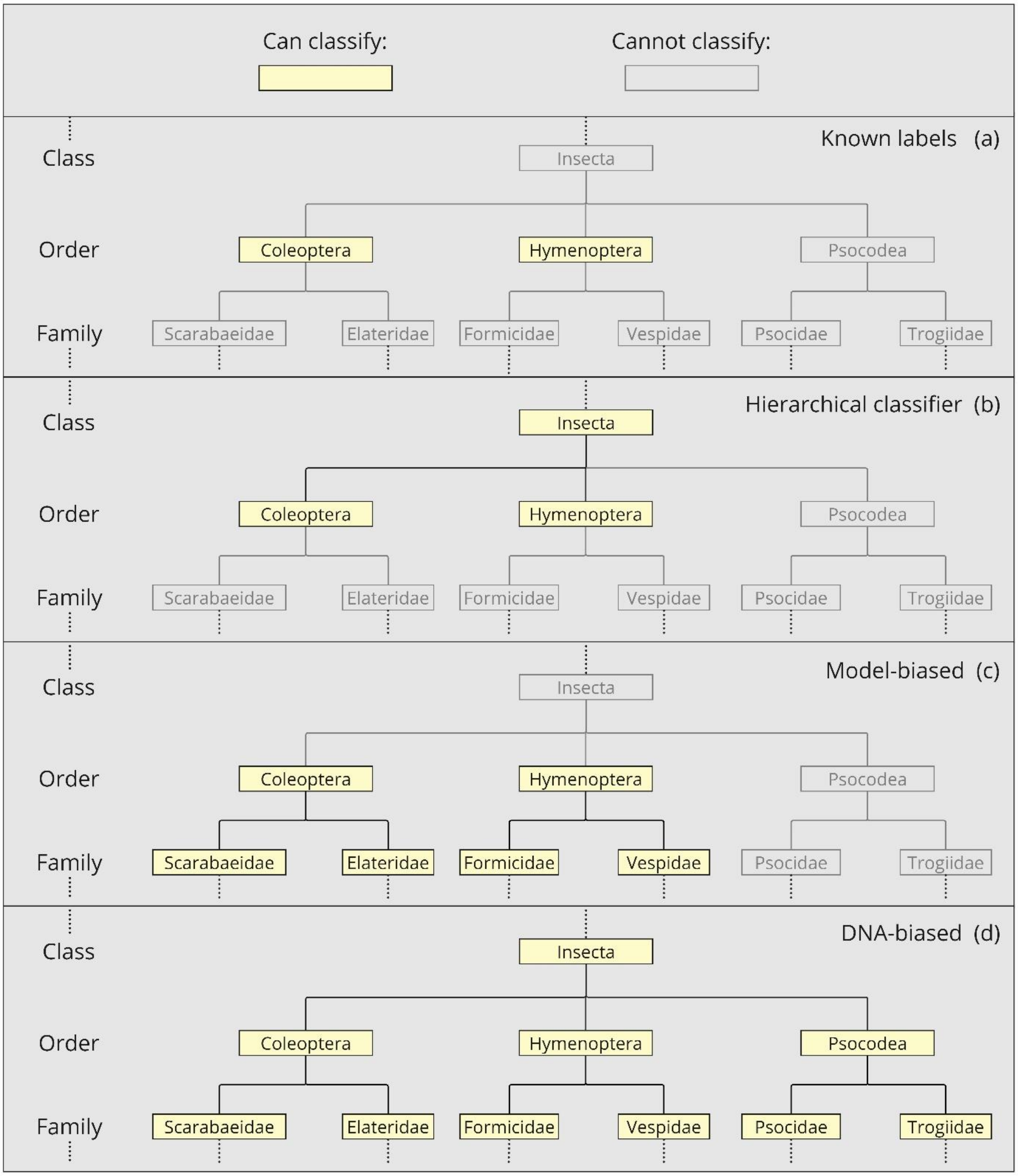


Figure S.5: Simplified example depicting the taxonomic scope of different classification methods. (a) The known classes of an example order level classifier. The base classifier can only infer these classes. (b) Hierarchical classification. The classifier can now infer its known classes and their supertaxa. (c) The model-biased taxonomic granularity improvement method. The classifier can now infer its known classes and their subtaxa. (d) The DNA-biased taxonomic granularity improvement method. The taxonomic scope is bounded only by the DNA reference database.

<u>Tables</u>

Table 1: Three examples of image-based hierarchical labels (a,b,c). (a) The hierarchical label for Blattodea with all taxonomic levels between phylum and order filled. (b) Some taxa do not have names for every taxonomic sub-level. In these cases, the name from the preceding, finer-grain level is used. In this example, Coleoptera does not belong to any superorder, so "Coleoptera" is used as a placeholder superorder name. (c) Classes with a taxonomic granularity above order-level used indeterminate ("indet.") labels at all remaining taxonomic levels. In this example, "Annelida indet." is used for all levels below phylum.

| Example | Phylum | Subphylum | Class | Subclass | Superorder | Order |
|---|---|---|---|---|---|---|
| (a) | Arthropoda | Hexapoda | Insecta | Pterygota | Polyneuroptera | Blattodea |
| (b) | Arthropoda | Hexapoda | Insecta | Pterygota | *Coleoptera* | Coleoptera |
| (c) | Annelida | *Annelida indet.* | *Annelida indet.* | *Annelida indet.* | *Annelida indet.* | *Annelida indet.* |

Table 2: Simplified example of a DNA multi-class hierarchical label for a sampling event. Our DNA hierarchical labels used 13 taxonomic levels from phylum to species, but for the sake of space only major taxonomic levels are shown here.

<table>
<tr><th>Phylum</th><th>Class</th><th>Order</th><th>Family</th><th>Genus</th><th>Species</th></tr>
<tr><td rowspan="4">Arthropoda</td><td rowspan="3">Insecta</td><td rowspan="2">Diptera</td><td rowspan="2">Culicidae</td><td>Aedes</td><td>A. vexans</td></tr>
<tr><td>Culex</td><td>C. salinarius</td></tr>
<tr><td>Orthoptera</td><td>Gryllidae</td><td>Gryllus</td><td>G. veletis</td></tr>
<tr><td>Arachnida</td><td>Araneae</td><td>Lycosidae</td><td>Schizocosa</td><td>S. humilis</td></tr>
</table>

Table S.3: Precision and recall values for each class when comparing sampling event assemblages detected by DNA metabarcoding against the ground truth image-based sampling event assemblages.

| Class | Precision | Recall |
|---|---|---|
| Annelida | 0.571 | 0.774 |
| Acari | 0.803 | 0.551 |
| Araneae | 0.942 | 0.852 |
| Opilioacarida | 1.000 | 0.000 |
| Opiliones | 0.750 | 0.120 |
| Collembola | 0.701 | 0.720 |
| Blattodea | 0.741 | 0.488 |
| Coleoptera | 0.934 | 0.752 |
| Diptera | 0.955 | 0.905 |
| Hemiptera | 0.542 | 0.640 |
| Hymenoptera | 0.984 | 0.529 |
| Lepidoptera | 0.229 | 0.905 |
| Orthoptera | 0.811 | 0.901 |
| Zygentoma | 1.000 | 0.000 |
| Isopoda | 0.909 | 0.667 |
| Myriapoda | 0.828 | 0.333 |
| Gastropoda | 0.718 | 0.549 |

Table S.4: Training dataset accuracy measurements for the four trained models.

| Model | Training accuracy |
|---|---|
| **Baseline** | 0.866 |
| **Fusion (DNA)** | 0.883 |
| **Fusion (zero-filled)** | 0.887 |
| **Fusion (“oracle”)** | 0.903 |

Table S.4: Specimen classifications from the DNA assemblage fusion model after applying taxonomic granularity refinement methods. The 'Label' column shows the identity of each classification. The 'DNA-biased' and 'Model-biased' count columns show the number of classifications for each label made by each respective granularity refinement method. The "Difference' column shows the absolute difference in the counts between the two methods. Subtotals are calculated for each taxonomic level.

| Level | Label | DNA-biased count | Model-biased count | Difference |
|---|---|---|---|---|
| **Species** | *Allonemobius maculatus* | 3 | 3 | - |
| | *Aphidius ervi* | 21 | 21 | - |
| | *Aporrectodea caliginosa* | 1 | 1 | - |
| | *Armadillidium nasatum* | 15 | 15 | - |
| | *Brachinus alternans* | 2 | 2 | - |
| | *Bradysia trivittata* | 4 | 4 | - |
| | *Deroceras laeve* | 6 | 6 | - |
| | *Diplocardia eiseni* | 1 | 1 | - |
| | *Diplocardia singularis* | 1 | 1 | - |
| | *Eleodes extricatus* | 3 | 3 | - |
| | *Eleodes tricostatus* | 7 | 7 | - |
| | *Eupithecia lariciata* | 1 | 1 | - |
| | *Homidia sinensis* | 1 | 1 | - |
| | *Leiobunum vittatum* | 40 | 10 | 30 |
| | *Lophopilio palpinalis* | 4 | 4 | - |
| | *Neoantistea magna* | 19 | 19 | - |
| | *Oxidus gracilis* | 3 | 3 | - |
| | *Oxypoda lacustris* | 1 | 1 | - |
| | *Parcoblatta bolliana* | 7 | 7 | - |
| | *Parcoblatta pensylvanica* | 3 | 3 | - |
| | *Parcoblatta uhleriana* | 1 | 1 | - |
| | *Pardosa distincta* | 64 | 64 | - |
| | *Pseudosinella octopunctata* | 46 | 46 | - |
| | *Scolopocryptops sexspinosus* | 3 | 3 | - |
| | *Stelidota geminata* | 6 | 6 | - |
| | *Stempellinella fimbriata* | 8 | 8 | - |
| | *Tinocallis ulmifolii* | 1 | 1 | - |
| | *Trachelipus rathkii* | 40 | 40 | - |
| | *Valenzuela flavidus* | 3 | 0 | 3 |
| | *Velarifictorus micado* | 4 | 4 | - |
| | **Subtotal** | **319** | **286** | **33** |

| Level | Label | DNA-biased count | Model-biased count | Difference |
|---|---|---|---|---|
| Genus | *Arion* | 39 | 39 | - |
| | *Balaustium* | 1 | 1 | - |
| | *Gryllus* | 38 | 38 | - |
| | *Lepidocyrtus* | 22 | 22 | - |
| | *Megaselia* | 21 | 21 | - |
| | *Myrmica* | 47 | 47 | - |
| | *Parcoblatta* | 637 | 11 | 626 |
| | *Photuris* | 4 | 4 | - |
| | *Schizocosa* | 17 | 17 | - |
| | *Solenopsis* | 272 | 272 | - |
| | *Stenomacrus* | 3 | 3 | - |
| | **Subtotal** | **1101** | **475** | **626** |
| Subfamily | Culicinae | 62 | 47 | 15 |
| | Lepidocyrtinae | 124 | 124 | - |
| | Lumbricinae | 27 | 27 | - |
| | Myrmicinae | 205 | 205 | - |
| | Pemphiginae | 2 | 2 | - |
| | **Subtotal** | **420** | **405** | **15** |
| Family | Bdellidae | 249 | 249 | - |
| | Entomobryidae | 2 | 2 | - |
| | Erythraeidae | 4 | 4 | - |
| | Eupodidae | 44 | 37 | 7 |
| | Isotomidae | 80 | 80 | - |
| | Julidae | 183 | 183 | - |
| | Lycosidae | 18 | 18 | - |
| | **Subtotal** | **580** | **573** | **7** |
| Superfamily | Blattoidea | 2 | 2 | - |
| | Grylloidea | 53 | 53 | - |
| | Oripodoidea | 6 | 6 | - |
| | Staphylinoidea | 1 | 1 | - |
| | **Subtotal** | **62** | **62** | **0** |
| Infraorder | Culicomorpha | 83 | 19 | 64 |
| | Ligiamorpha | 3 | 3 | - |
| | Muscomorpha | 24 | 15 | 9 |
| | Neolepidoptera | 2 | 2 | - |
| | **Subtotal** | **112** | **39** | **73** |
| Suborder | Apocrita | 178 | 178 | - |
| | Araneomorphae | 512 | 297 | 215 |
| | Ensifera | 26 | 26 | - |

| Level | Label | DNA-biased count | Model-biased count | Difference |
|---|---|---|---|---|
| **Suborder** | Entomobryomorpha | 116 | 116 | - |
| | Nematocera | 309 | 139 | 170 |
| | Polyphaga | 3 | 3 | - |
| | Prostigmata | 2 | 2 | - |
| | Psocomorpha | 3 | 0 | 3 |
| | Symphypleona | 10 | 10 | - |
| | **Subtotal** | **1159** | **771** | **388** |
| **Order** | Araneae | 86 | 88 | 2 |
| | Coleoptera | 375 | 466 | 91 |
| | Collembola | 68 | 130 | 62 |
| | Diptera | 446 | 655 | 209 |
| | Hemiptera | 9 | 26 | 17 |
| | Lepidoptera | 1 | 1 | - |
| | Orthoptera | 13 | 25 | 12 |
| | Sarcoptiformes | 46 | 46 | - |
| | Stylommatophora | 164 | 164 | - |
| | Blattodea | 0 | 2 | 2 |
| | Hymenoptera | 0 | 853 | 853 |
| | Isopoda | 0 | 18 | 18 |
| | Opilioacarida | 0 | 40 | 40 |
| | Opiliones | 0 | 116 | 116 |
| | Zygentoma | 0 | 18 | 18 |
| | **Subtotal** | **1208** | **2648** | **1440** |
| **Superorder** | Acariformes | 86 | 78 | 8 |
| | Holometabola | 269 | 0 | 269 |
| | **Subtotal** | **355** | **78** | **277** |
| **Subclass** | Acari | 6 | 242 | 236 |
| | Pterygota | 144 | 0 | 144 |
| | **Subtotal** | **150** | **242** | **380** |
| **Class** | Arachnida | 85 | 0 | 85 |
| | Diplopoda | 4 | 4 | - |
| | Gastropoda | 6 | 6 | - |
| | Insecta | 20 | 0 | 20 |
| | **Subtotal** | **115** | **10** | **105** |
| **Subphylum** | Myriapoda | 0 | 27 | 27 |
| | **Subtotal** | **0** | **27** | **27** |
| **Phyum** | Annelida | 1 | 1 | - |
| | Arthropoda | 35 | 0 | 35 |
| | **Subtotal** | **36** | **1** | **35** |
